# Mammalian NSUN2 introduces 5-methylcytidines into mitochondrial tRNAs

**DOI:** 10.1101/624932

**Authors:** Saori Shinoda, Sho Kitagawa, Shinichi Nakagawa, Fan-Yan Wei, Kazuhito Tomizawa, Kimi Araki, Masatake Araki, Takeo Suzuki, Tsutomu Suzuki

**Affiliations:** Department of Chemistry and Biotechnology, Graduate School of Engineering, the University of Tokyo, 7-3-1 Hongo, Bunkyo-ku, Tokyo 113-8656, Japan; Faculty of Pharmaceutical Sciences, Hokkaido University, Hokkaido 060-0812, Japan; Department of Molecular Physiology, Faculty of Life Sciences, Kumamoto University, Kumamoto 860-8556, Japan; Research for Embryonic Science and Technology (PRESTO), Japan Science and Technology Agency (JST), Kawaguchi, Japan; Institute of Resource Development and Analysis, Kumamoto University, Kumamoto 860-0811, Japan

**Author notes:** These authors contributed equally to this work (Saori Shinoda, Sho Kitagawa).

## Abstract

Post-transcriptional modifications in mitochondrial tRNAs (mt-tRNAs) play critical roles in mitochondrial protein synthesis, which produces respiratory chain complexes. In this study, we used mass spectrometric analysis to map 5-methylcytidine (m^5^C) at positions 48–50 in eight mouse and six human mt-tRNAs. We also confirmed the absence of m^5^C in mt-tRNAs isolated from *Nsun2* knockout (KO) mice, as well as from *NSUN2* KO human culture cells. In addition, we successfully reconstituted m^5^C at positions 48–50 of mt-tRNA *in vitro* with NSUN2 protein in the presence of *S*-adenosylmethionine (SAM). Although NSUN2 is predominantly localized to the nucleus and introduces m^5^C into cytoplasmic tRNAs and mRNAs, structured illumination microscopy (SIM) clearly revealed NSUN2 foci inside mitochondria. These observations provide novel insights into the role of NSUN2 in the physiology and pathology of mitochondrial functions.

## INTRODUCTION

The mitochondrion is a eukaryotic organelle that performs aerobic respiration through the electron transport chain and oxidative phosphorylation, thereby generating chemical energy in the form of ATP. Mitochondria have their own genome (mitochondrial DNA: mt-DNA) and a devoted gene expression system, consisting of transcriptional and translational machinery, that produces 13 subunits of respiratory chain complexes encoded in mt-DNA (1-3).

In mammals, the mitochondrial translational apparatus requires 22 species of mitochondrial (mt-)tRNAs encoded in mt-DNA. Long polycistronic precursor RNAs are transcribed from both strands of mt-DNA and processed into rRNAs, mRNAs, and tRNAs. Because mt-tRNAs lie between mRNAs and rRNAs in the precursor RNAs, mt-tRNA processing results in separation of mRNAs and rRNAs (4). During the process of maturation, mt-tRNAs undergo various post-transcriptional modifications (3). We previously identified 15 species of modified nucleosides at 118 positions in all 22 mt-tRNAs of bovine mitochondria (5). mt-tRNAs are more heavily modified than mt-rRNAs, which are modified at only ten positions in the 12S and 16S rRNAs (6). In particular, a wide variety of modifications are present in the anticodon region of tRNAs (3), and these modifications play critical roles in the precise decoding of genetic codes (7).

The functional and physiological importance of mt-tRNA modifications is highlighted by the observation that mt-tRNAs are hypomodified in the cells of patients with mitochondrial diseases (3,8). We previously reported that 5-taurinomethyluridine (τm^5^U) and its 2-thiouridine derivative (τm^5^s^2^U) at the wobble position (position 34) of the anticodon (9) are not formed in mutant mt-tRNAs^Leu(UUR)^isolated from patients with MELAS (mitochondrial encephalopathy, lactic acidosis, and stroke-like syndrome) or in mutant mt-tRNA^Lys^isolated from patients with MERRF (myoclonus epilepsy with ragged-red fibers), respectively (3,8). The absence of these taurine modifications results in defective mitochondrial translation, leading to mitochondrial dysfunction. Supporting these findings, several pathogenic mutations associated with mitochondrial disorders have been reported in the human genes *MTO1, GTPBP3*, and *MTU1*, which encode enzymes involved in τm^5^s^2^U biogenesis (10,11).

5-formylcytidine (f^5^C), another unique modification found at the wobble position of mt-tRNA^Met^(12), plays a critical role in deciphering the AUA codon as Met. We reported that NSUN3 (13) and ALKBH1 (14) are responsible for f^5^C34 biogenesis, and identified two pathogenic point mutations in mt-tRNA^Met^that impair f^5^C34 formation (13). *N*^6^-threonylcarbamoyladenosine (t^6^A) occurs at position 37, 3’-adjacent to the anticodon of five mt-tRNA species (5). We also showed that YRDC and OSGEPL1 are responsible for t^6^A37 formation, and demonstrated that *OSGEPL1*-knockout cells exhibit mitochondrial dysfunction (15). Moreover, we found that levels of t^6^A37 are reduced in mutant mt-tRNA isolated from the cells of patients with MERRF-like symptoms (15).

Modifications in the anticodon region of mt-tRNAs are required for mitochondrial translation, and the absence of these modifications has pathological consequences. By contrast, the physiological importance of modifications in the tRNA body region of tRNAs remains elusive. 5-methylcytidine (m^5^C) is a common modification in tRNA body regions (16). In eukaryotes, m^5^C is present in a wide variety of RNA species, including tRNA, rRNA, mRNA, and viral RNA (17-20) (Figure 1A). m^5^C is introduced by the NOL1/NOP2/SUN domain (NSUN) family of enzymes, as well as by DNMT2 (21). Cytoplasmic tRNAs (ct-tRNAs) contain m^5^C at positions 34, 38, 40, 48–50, and 72. m^5^C38 and m^5^C72 are introduced by DNMT2 (22) and NSUN6 (23), respectively, whereas m^5^Cs at the other positions are introduced by NSUN2 (24-27). Loss of m^5^C in double-knockout mice of *Nsun2* and *Dnmt2* induces tRNA cleavage mediated by angiogenin to generate 5’ tRNA fragments (28,29) and decreases the steady-state levels of ct-tRNAs (27). In addition, these tRNA fragments inhibit cap-dependent translation (30,31). Loss-of-function mutations in the human *NSUN2* gene are associated with autosomal recessive intellectual disabilities such as Dubowitz syndrome (26,32), and *Nsun2* KO mice exhibit weight loss, microcephaly, male infertility, and partial alopecia (33). Despite our knowledge of the importance of m^5^C in ct-tRNAs, little is known about the biogenesis and function of m^5^C in mt-tRNAs. In several mt-tRNA species, m^5^C is present at positions 48–50 in the extra loop (Figures 1B and S1) (5,34). We previously speculated that NSUN2 also participates in m^5^C formation of mt-tRNAs (3), even though NSUN2 is predominantly localized to nucleus (32).

**Figure 1.**
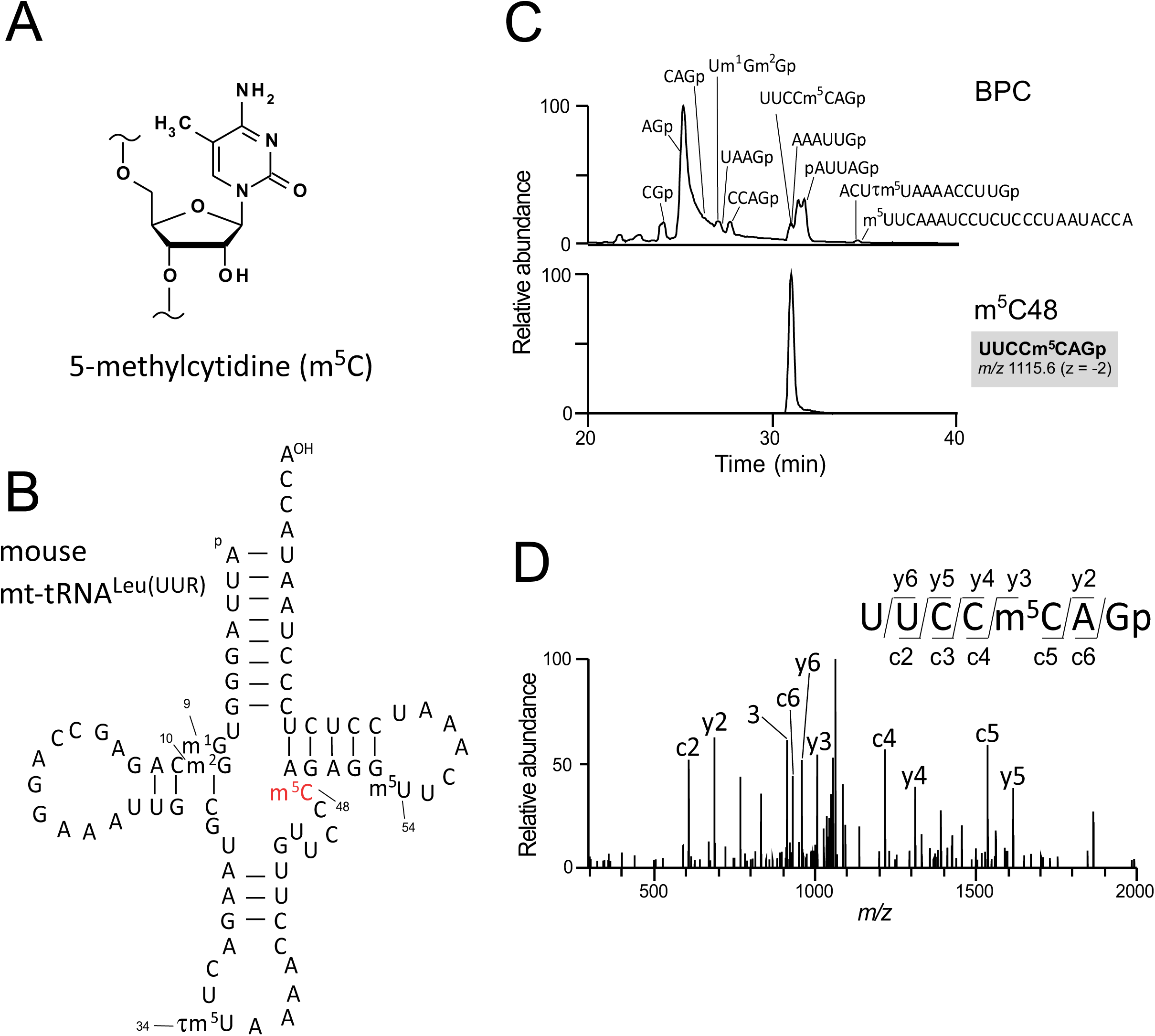
mt-tRNAs contain m^5^C in the extra loop. (A) Chemical structure of 5-methylcytidine (m^5^C). (B) Secondary structure of mouse mitochondrial tRNA^Leu(UUR)^with modified nucleosides: 1-methylguanosine (m^1^G), *N*^*2*^-methylguanosine (m^2^G), 5-taurinomehtyluridine (τm^5^U), 5-methylcytidine (m^5^C), and 5-methyluridine (m^5^U). Pseudouridine is not shown. (C) RNA fragment analysis of mouse mt-tRNA^Leu(UUR)^ digested with RNase T_1_. Assigned fragments are indicated on the base peak chromatogram (BPC) in the upper panel. ‘p’ represents a terminal phosphate group. An extracted-ion chromatogram (XIC) for the doubly charged negative ion of an m^5^C-containing fragment (UUCCm^5^CAGp *m/z* 1115.64) is shown in the lower panel. (D) Collision-induced dissociation (CID) spectrum of the m^5^C-containing fragment shown in (C) used to determine the m^5^C site. The doubly charged negative ion of the fragment UUCCm^5^CAGp (*m/z* 1115.64) was used as the precursor ion. Assigned c-and y-series product ions are indicated in the spectrum.

Here, we report precise mapping of m^5^C in mt-tRNAs, and present clear evidence that NSUN2 is partially localized to mitochondria and introduces m^5^C in mt-tRNAs.

## MATERIALS AND METHODS

### Cell culture

HeLa and HEK293T cells, and mouse embryonic fibroblast (MEFs) were cultured at 37°C in an atmosphere containing 5% CO2 in Dulbecco’s modified Eagle’s medium (DMEM, Sigma-Aldrich) supplemented with 10% fetal bovine serum (FBS, Gibco) and 1% penicillin–streptomycin (PS, FUJIFILM Wako Pure Chemical Corporation).

### Animals

*Nsun2*^-/-^mice (gene trap clone ID 21-B114 in the EGTC database, http://egtc.jp/) (35) were provided by the Center for Animal Resources and Development (CARD) at Kumamoto University. Animals were housed at 25°C with 12h light and 12h dark cycles. To generate MEFs, male and female *Nsun2*^+/-^mice were mated, and embryos were dissected out at day 14 of gestation. Embryo head and liver were removed for genotyping, the remaining tissues were treated with trypsin for 10 min at 37°C. Primary MEFs were seeded in 10 mm dishes and stored at passage number 2. All animal procedures were approved by the Animal Ethics Committee of Kumamoto University (Approval ID: A29-016R3).

### Construction of *NSUN2* KO cell lines

Oligonucleotide sequences used for gene editing are listed in Table S1. *NSUN2* KO HEK293T cell lines were constructed using the CRISPR-Cas9 system as described previously (13,36). In brief, sense and antisense oligonucleotides for a single guide RNA (sgRNA) were cloned into pX330 (Addgene plasmid #42230) (37). HEK293T cells seeded on 24-well plates were co-transfected with 300 ng pX330 containing the sgRNA sequence, 100 ng pEGFP-N1 (Clontech), and 100 ng modified pLL3.7 vector containing a puromycin resistance gene. Transfections were performed using FuGENE HD (Promega). The following day, the cells were sparsely seeded in a 100 mm dish, and transfectants were selected with 1 μg/mL puromycin. Transfection efficiency was assessed by monitoring EGFP fluorescence. A few days after the transfection, several colonies were isolated and grown for an additional week. The sequence of the targeted region in each selected clone was confirmed by direct sequencing of genomic PCR products. PCR primer sequences are provided in Table S1.

### RNA preparation, tRNA isolation, and mass spectrometry

Total RNA from culture cells was prepared by a standard acid guanidium–phenol– chloroform (AGPC) method (38). Individual mt-tRNAs were isolated by reciprocal circulation chromatography (RCC) (39). DNA probe sequences complementary to each mt-tRNA are listed in Table S1. Capillary LC–nano-ESI–mass spectrometry (RNA-MS) of mt-tRNA fragments digested by RNase T_1_ was performed as described (5,40).

### Northern blotting

Total RNA (2 μg) from mouse liver was separated by 10% denaturing polyacrylamide gel electrophoresis (PAGE) and blotted onto a nylon membrane (Amersham Hybond N^+^; GE Healthcare) using a Transblot Turbo apparatus (Bio-Rad). The blotted RNA was crosslinked to the membrane by two rounds of irradiation with UV light (254 nm, 120 mJ/cm^2^ for one round; CL-1000, UVP). DNA probes were 5’-phosphorylated with T4 polynucleotide kinase (PNK, TOYOBO) and [γ-^32^P] ATP (PerkinElmer). Northern blotting of tRNAs was performed using PerfectHyb (Toyobo) at 48–55°C with 4 pmol of labeled DNA probes specific to tRNAs and 3 pmol of a labeled DNA probe (mixed with 9 pmol of non-labeled probe) specific for 5S rRNA. The membrane was washed six times with 1× SSC (150 mM NaCl and 15 mM sodium citrate, adjusted to pH 7.0 with citric acid), and exposed to an imaging plate (BAS-MS2040, Fujifilm). Radioactivity was visualized on an FLA-7000 imaging system (Fujifilm). To quantify the steady-state level of the target tRNA, the radioactivity of the tRNA band was normalized against the 5S rRNA band.

### Expression and isolation of NSUN2

Human *NSUN2* cDNA was reverse-transcribed from total RNA of HeLa cells, amplified using primers listed in Table S1, and cloned into the entry vector pENTR/D-TOPO (Invitrogen). The gene cassette containing the entry clone was then transferred to a modified pDEST12.2 (Invitrogen) harboring a C-terminal FLAG tag.

For transient expression of NSUN2, HEK293T cells (4 × 10^6^) in 100 mm dishes were transfected with 10 µg pDEST12.2-NSUN2-FLAG using Lipofectamine 2000 (Invitrogen), and cultured for 48–72 h at 37°C in 5% CO_2_. The cells were suspended with 1 mL of lysis buffer [50 mM HEPES-KOH (pH 7.9), 250 mM KCl, 2 mM MgCl_2_, 0.5 mM DTT, 0.5% Triton-X100, and 1 × Complete EDTA-free protease inhibitor cocktail (Roche Life Science)] and lysed by sonication. The lysate was centrifuged twice at 20,000 × *g* for 20 min at 4°C to remove cell debris. The supernatant was immunoprecipitated with 50 µL of a 50% slurry of anti-FLAG M2 agarose beads (Sigma-Aldrich). The beads were washed three times with 500 µL lysis buffer, and NSUN2 was eluted from the beads at 4°C for 3 h in elution buffer [150 mM KCl, 10 mM Tris-HCl (pH 8.0), 20% Glycerol, 0.1 mM DTT, and 200 µg/mL DYKDDDDK peptide (FUJIFILM Wako Pure Chemical Corporation)]. Isolated NSUN2 was quantified based on the intensity of the corresponding band in an SDS-PAGE gel stained with SYPRO Ruby (Thermo Fisher Scientific), using BSA as a standard.

### *In vitro* m^5^C formation using NSUN2

Substrate mt-tRNA^Ser(AGY)^was transcribed *in vitro* with T7 RNA polymerase as described (41). Oligonucleotide sequences for preparation of template DNA are listed in Table S1. The transcript was run on a 10% denaturing PAGE gel, and the intact tRNA band was excised from the gel, followed by tRNA extraction. *In vitro* methylation was performed at 37°C for 2 h in 50 μL of a reaction mixture consisting of 20 mM HEPES-KOH (pH 8.0), 5 mM MgCl_2_, 100 mM KCl, 1 mM DTT, 0.5 μM mt-tRNA^Ser(AGY)^ transcript, 0.5 μM NSUN2, and 1 mM *S*-adenosylmethionine (SAM). The tRNA was extracted with phenol and precipitated with ethanol. A 1.5 pmol aliquot of the tRNA was digested with RNase T_1_ and subjected to LC/MS analysis to detect m^5^C-containing fragments.

### Fluorescence microscopy

HeLa cells grown on poly-L-lysine–coated coverslips were incubated in 5% CO_2_ at 37°C for 1 h with 5 µM MitoTracker Red CMXRos (Molecular Probes) in DMEM. The cells were washed with phosphate-buffered saline (PBS), fixed with 3.7% formaldehyde in PBS for 10 min at room temperature, permeabilized with 0.5% Triton X-100 in PBS for 10 min at room temperature, blocked with 2% FBS in PBS for 30 min at room temperature, and incubated at room temperature for 1 h with anti-NSUN2 antibody (1:500; SIGMA HPA037896) or anti-TFAM antibody (1:500; Abnova E7031) diluted in Can Get Signal solution A (Toyobo). After three washes in PBS, the cells were incubated at room temperature for 1 h with Cy2/Cy3-conjugated anti-mouse or anti-rabbit secondary antibody (1:50; Millipore AP182C). After three washes in PBS, the cells were mounted in 97% 2,2’-thiodiethanol containing 2% 1,4-diazabicyclo[2.2.2]octane. The coverslips were observed using a confocal microscope (FV1000; Olympus) with a 60 × oil immersion objective lens (1.42 BNA; Olympus). Images were acquired using the FluoView software (Olympus) and processed using ImageJ (42). For 5-bromouridine (BrU) labeling, HeLa cells on coverslips were incubated with 2.5 mM BrU for 20 min prior to fixation. Anti-BrU antibody (1:50; Roche 11170376001) and anti-NSUN2 antibody (1:500; SIGMA HPA037896) were used to detect BrU-labeled RNA and NSUN2, respectively. Secondary antibodies conjugated with Alexa fluor 555 (1:1000; Invitrogen A-21422) and Alexa fluor 488 (1:1000; Invitrogen A-11008) were subsequently used respectively. The coverslips were observed using a confocal microscope (FV3000; Olympus).

### Super-resolution structured illumination microscopy (SR-SIM)

Super-resolution imaging was performed on an ELYRA PS.1 microscope (Carl Zeiss) equipped with a alpha Plan-Apochromat 100×/1.46 Oil DIC M27 ELYRA objective and EM-CCD camera (12.1 × 12.1 µm^2^ field consisting of 256 × 256 pixels) as previously reported (43). To observe mitochondrial foci of NSUN2, 20 Z-series images were obtained at 100 nm intervals with the 561 nm and 481 nm lasers, and SIM images were calculated using default settings with theoretically predicted point-spread function parameters. For channel alignment, 0.2 μm multicolored beads (TetraSpec Microspheres, Thermo Fisher) immobilized on a sample coverslip were imaged using the same acquisition settings, and the resultant images were used to correct for chromatic shifts between color channels. Images were processed using the ZEN software (Carl Zeiss).

### Pulse labeling of mitochondrial protein synthesis

The pulse-labeling experiment was performed essentially as described (13). WT and *NSUN2* KO HEK293T cells (2.0 × 10^6^) were cultured at 37°C in 5% CO_2_ for 10 min in methionine-, glutamine-, and cysteine-free DMEM (Gibco) supplemented with 2 mM L-glutamine, 10% FBS, and 50 μg/mL emetine (to inhibit cytoplasmic protein synthesis). The cells were then supplemented with 7.4 MBq of [^35^S] methionine and [^35^S] cysteine (EXPRE^35^S^35^S Protein Labeling Mix, [^35^S]-, PerkinElmer) and incubated for 1 h to specifically label newly synthesized mitochondrial proteins. Cell lysates (50 μg total proteins) were resolved by tricine–SDS-PAGE (16.5%), and the gel was CBB stained and dried on a gel drier (AE-3750 RapiDry, ATTO). Radiolabeled mitochondrial protein products were visualized using an imaging plate (BAS-MS2040, Fujifilm) on an FLA-7000 imaging system.

### Steady-state levels of subunit proteins in respiratory chain complexes

Mitochondria were isolated from WT and *NSUN2* KO HEK293T cells (1 × 10^7^) using the Mitochondria Isolation Kit (Miltenyi Biotec). Steady-state levels of subunit proteins of mitochondrial respiratory chain complexes were analyzed by immunoblotting with Total OXPHOS Rodent WB Antibody Cocktail (1:250, ab110413, Abcam) and anti-mt-ND5 antibody (1:100, ab92624, Abcam). HRP-conjugated anti-mouse IgG (1:20,000, 715-035-150, Jackson ImmunoResearch) or HRP-conjugated anti-rabbit IgG (1:20,000, 715-035-152, Jackson ImmunoResearch) was used as the secondary antibody.

### Resazurin-based cell proliferation assay

WT, *NSUN2* KO, and *NSUN3* KO HEK293T cells were seeded on 96-well plates (1.0 × 10^5^cells/well) and cultured in glucose-free DMEM (Gibco) containing 10% FBS, 1% PS, 1 mM sodium pyruvate, and 4% glucose or in glucose-free DMEM (Gibco) containing 10% FBS, 1% PS, 1 mM sodium pyruvate, and 4% galactose. On each day, 1/10 volume of 1 mM resazurin solution in PBS was added to each well and incubated for 3 h. Absorbance was measured at 570 and 600 nm on a microplate reader (SpectraMax Paradigm, Molecular Devices). The reduction rate of resazurin was calculated using the following equation:

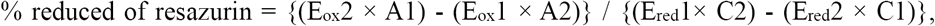

where

E_ox_1 = molar extinction coefficient (E) of oxidized (ox) resazurin at 570 nm = 80586,

E_ox_2 = E of oxidized resazurin at 600 nm = 117216,

E_red_1 = E of reduced (red) resazurin at 570 nm = 155677,

E_red_2 = E of reduced resazurin at 600 nm = 14652,

A1 = absorbance of measured well at 570 nm,

A2 = absorbance of measured well at 600 nm,

C1 = absorbance of blank well (medium and resazurin only) at 570 nm, and

C2 = absorbance of blank well at 600 nm.

## RESULTS

### Mass spectrometric m^5^C mapping of mouse and human mt-tRNAs

To map m^5^C, we isolated eight species of mouse mt-tRNAs bearing C at positions 48–50 [for Glu, His, Met, Asn, Leu(UUR), Leu(CUN), Ser(AGY), and Tyr] from mouse liver using reciprocal circulating chromatography (RCC) (39). Purified tRNAs were digested by RNase T_1_ and subjected to capillary LC-nano-ESI-mass spectrometry (RNA-MS) to analyze post-transcriptional modifications (15,40,44,45). Each RNA fragment was assigned by comparing observed *m/z* values with the calculated values (Figure 1C). The sequence of each fragment was further analyzed by collision-induced dissociation (CID) to determine the precise position of each modification (Figure 1D). We mapped m^5^C at eight positions in the extra loop of the eight tRNA species: at position 48 of five mt-tRNAs [His, Met, Leu(UUR), Asn, and Tyr], and at position 49 of three mt-tRNAs [Glu, Leu(CUN), and Ser(AGY)] (Figures S1 and S2).

In a similar manner, we isolated six species of human mt-tRNAs bearing C at positions 48–49 [for Glu, Phe, His, Leu(UUR), Ser(AGY), and Tyr] from HEK293T cells. RNA-MS analyses detected m^5^C at eight positions in the six tRNA species: at position 48 of five mt-tRNAs [Phe, His, Leu(UUR), Ser(AGY), and Tyr], at position 49 of two mt-tRNAs [Glu and Ser(AGY)], and position 50 of mt-tRNA^Ser(AGY)^(Figures S1 and S3).

### *NSUN2* is essential for m^5^C formation in mammalian mt-tRNA

Given that NSUN2 introduces m^5^C into the extra loop of ct-tRNAs (24-27), we speculated that NSUN2 is also responsible for m^5^C formation in mammalian mt-tRNAs (3). To test this hypothesis, we analyzed tRNA modification status in an *Nsun2*-null mouse constructed by gene trapping (Figure 2A). We isolated ct-tRNAs as well as mt-tRNAs from the livers of *Nsun2*^-/-^mice, and analyzed m^5^C48/49/50 state by RNA-MS. As observed previously (27), m^5^C was not present in ct-tRNA^Gly^isolated from liver from *Nsun2*^-/-^mouse (Figure 2B). We also found that m^5^C was completely absent in eight mt-tRNAs isolated from *Nsun2*^-/-^mouse livers (Figures 2C and S2).

**Figure 2.**
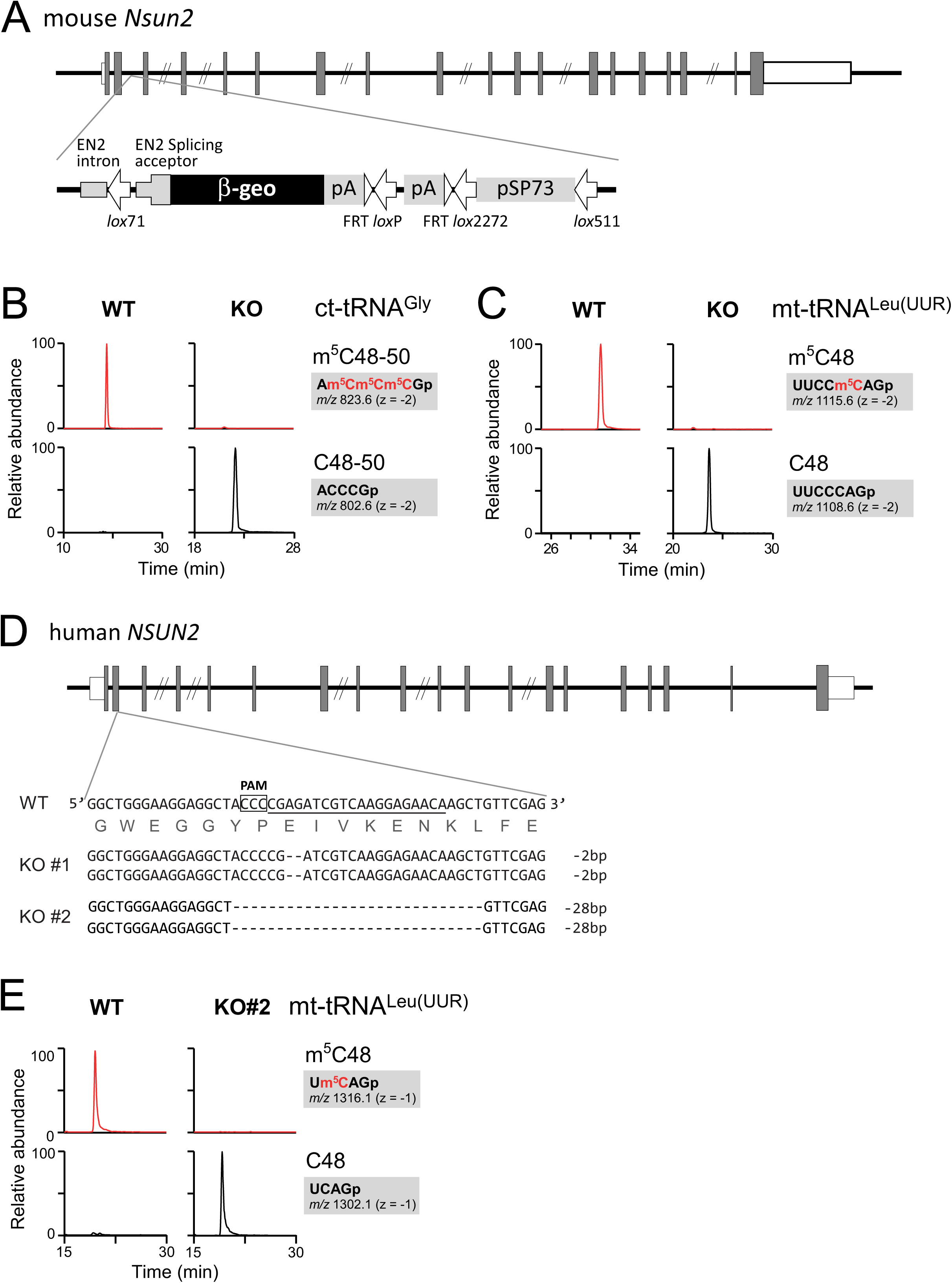
*NSUN2* is responsible for m^5^C formation in mt-tRNAs. (A) Schematic depiction of the mouse *Nsun2* gene with the insertion site of the gene trap cassette containing the β-geo marker. Shaded boxes, open boxes, and lines indicate coding regions, untranslated regions of exons, and introns, respectively. (B, C) XICs of RNase T_1_-digested fragments of mouse ct-tRNA^Gly^ (B) and mt-tRNA^Leu(UUR)^(C) containing m^5^C (top) and C (bottom) isolated from WT and *Nsun2*^-/-^(KO) mouse liver. (D) Schematic depiction of the human *NSUN2* gene with mutation sites introduced by the CRISPR-Cas9 system. The target sequence of the single guide RNA (sgRNA) is underlined. The protospacer adjacent motif (PAM) sequence is boxed. Deletions are represented by dashed lines. (E) XICs of RNase T_1_-digested fragments of human mt-tRNA^Leu(UUR)^ containing m^5^C (top) and C (bottom) isolated from WT and *NSUN2* KO #2 HEK293T cells.

For human culture cells, we knocked out the *NSUN2* gene in HEK293T cells using CRISPR/Cas9, yielding two KO cell lines (KO #1 and KO #2) in which both alleles harbored a homozygous frameshift deletion (Figure 2D). We then isolated six mt-tRNAs from *NSUN2* KO cells and subjected them to RNA-MS. As expected, no m^5^C was detected in any of the isolated tRNAs (Figure S3), demonstrating that *NSUN2* is required for the formation of m^5^C in mt-tRNAs.

### *In vitro* reconstitution of m^5^C in mt-tRNA

We next performed *in vitro* reconstitution of m^5^C in mt-tRNA using recombinant NSUN2. For these experiments, human NSUN2 fused with a C-terminal FLAG tag was expressed in HEK293T cells and immunoprecipitated with anti-FLAG antibody. As a substrate, we prepared an *in vitro* transcript of human mt-tRNA^Ser(AGY)^, which was incubated with human NSUN2 and *S*-adenosylmethionine (SAM). After the reaction, the tRNA substrate was digested with RNase T_1_ and subjected to RNA-MS. We clearly detected RNA fragments bearing mono-, di-, and trimethylations (Figure 3A). CID analysis revealed that trimethylation occurred at cytidines at positions 48–50 (Figure 3B). This result is consistent with our observation that human mt-tRNA^Ser(AGY)^had three m^5^C residues at positions 48–50 (Figures S1 and S3).

**Figure 3.**
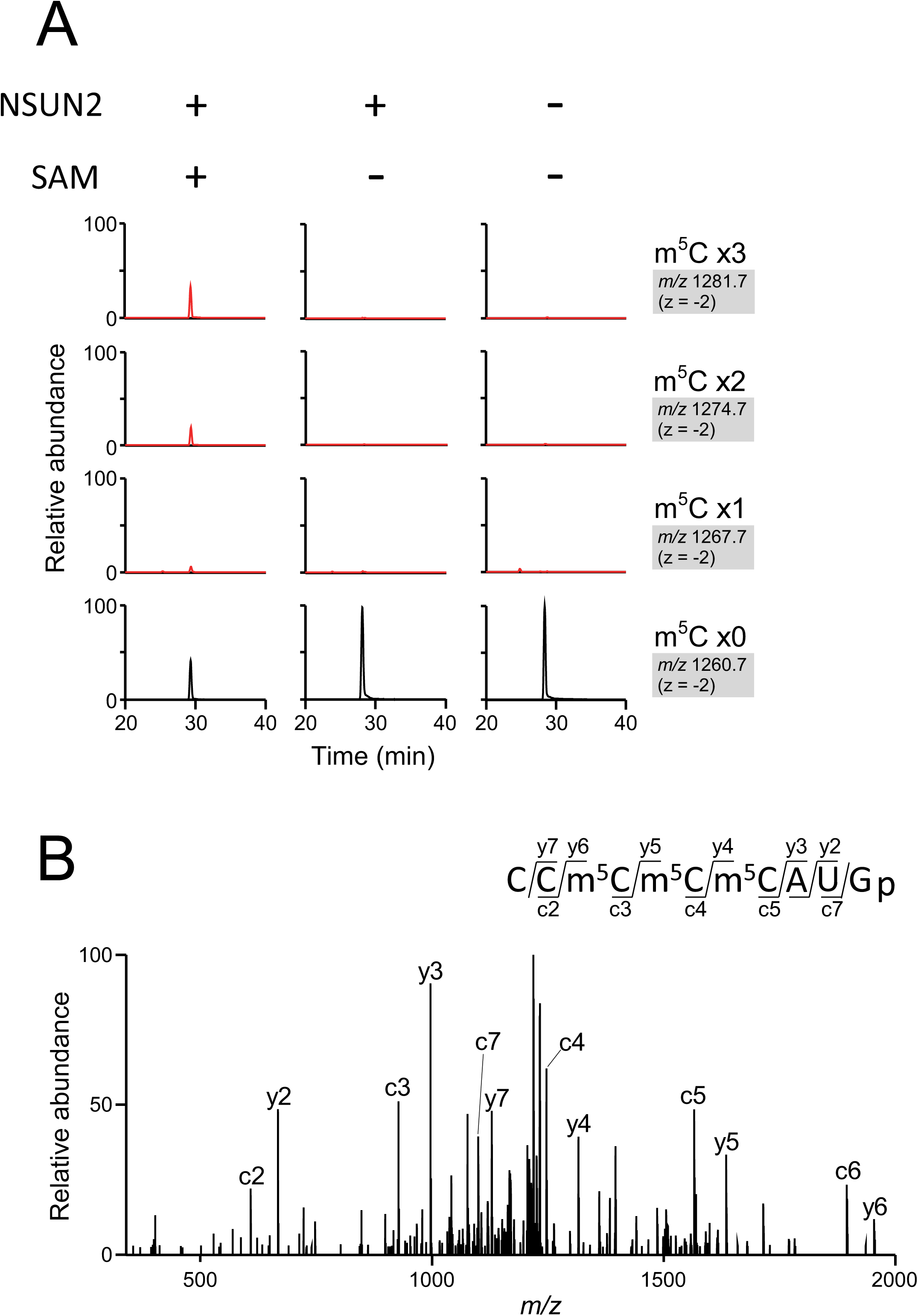
*In vitro* reconstitution of m^5^C on mt-tRNA. (A) NSUN2-mediated m^5^C formation on the human mt-tRNA^Ser(AGY)^transcript in the presence or absence of recombinant NSUN2 and SAM. XICs of RNase T_1_-digested fragments (CCCCCAUGp) of human mt-tRNA^Ser (AGY)^containing three (top panels), two (second panels), one (third panels), or zero (bottom panels) methyl groups. *m/z* value with the charge state for each fragment is shown on the right. (B) CID spectrum of the trimethylated fragment in (A). The precursor ion for CID is *m/z* 1281.69. Assigned c-and y-series product ions are indicated in the spectrum.

### Subcellular localization of endogenous NSUN2

NSUN2 is predominantly localized to the nucleus (32). To explore the possibility that some of the enzyme is targeted to mitochondria, we immunostained endogenous NSUN2 in HeLa cells with anti-NSUN2 antibody and visualized the protein by confocal microscopy. Most NSUN2 signal was observed in nucleus (Figure 4A), as reported previously (32), but some of the signal was dispersed in the cytoplasm and partially overlapped with mitochondria (Figure 4A). We further investigated cytoplasmic NSUN2 signals using super-resolution structured illumination microscopy (SR-SIM) (46). The increase in both the lateral (*x*-*y* axis) and axial (*z*-axis) resolution of SR-SIM enabled visualization of the cytoplasmic NSUN2 signal inside mitochondria (Figure 4B). Furthermore, signals of mitochondrial NSUN2 partially overlapped with those of BrU-labeled RNAs (Figure 4C) which represent newly-synthesized transcripts in mitochondria, whereas mitochondrial NSUN2s were observed as signals that reside next to TFAM-stained signals that co-localize with mt-DNA (47) (Figure 4D). This observation implies that NSUN2-mediated m^5^C formation takes place co-transcriptionally in the precursor transcripts. Taken together, these observations show that while NSUN2 is predominantly localized to the nucleus, it is also present in mitochondria, consistent with a role in forming m^5^C formation at positions 48–50 of mt-tRNAs near the site of transcription.

**Figure 4.**
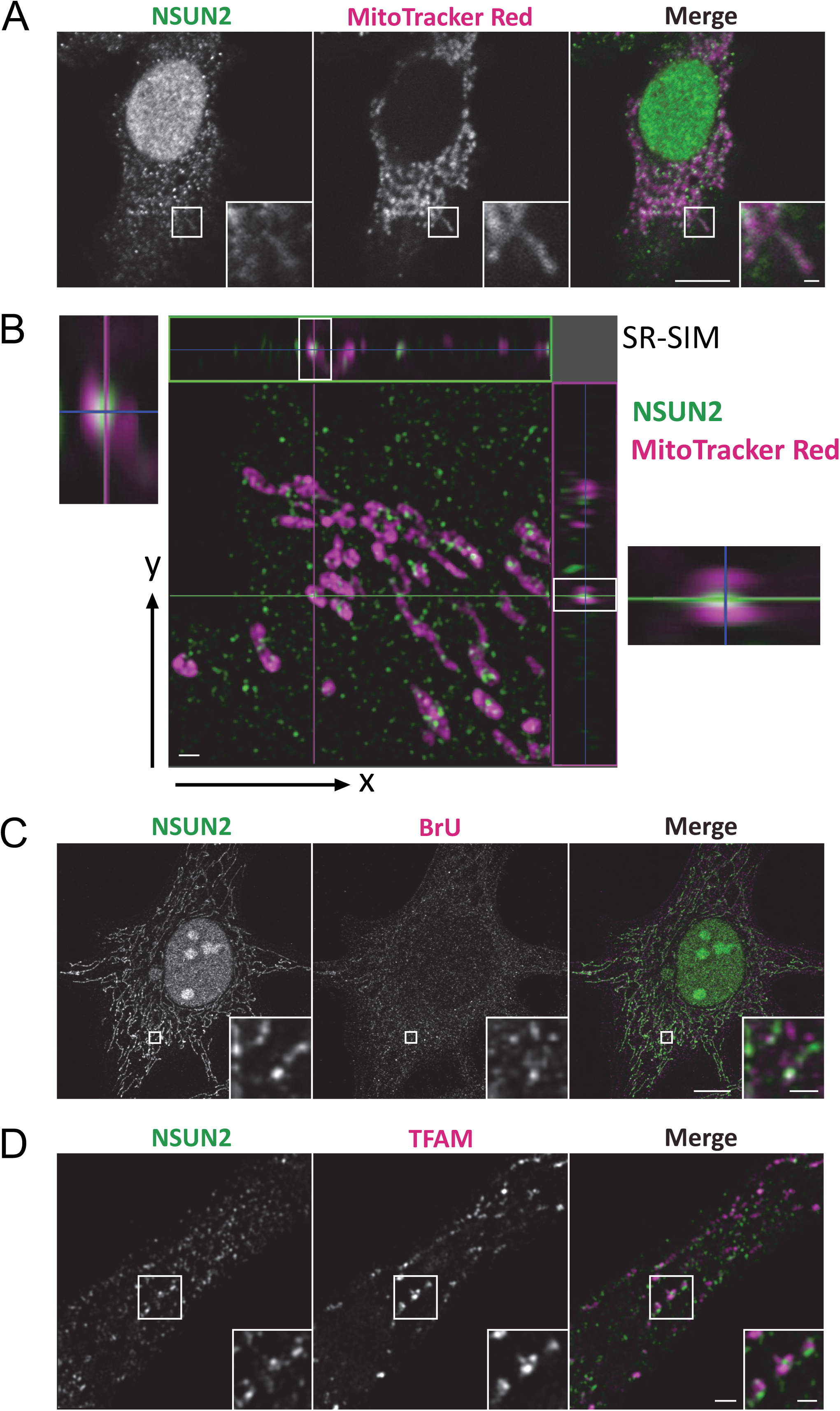
Subcellular localization of endogenous human NSUN2. (A, B) Subcellular localization of human NSUN2 in HeLa cells showing NSUN2 (green) and MitoTracker Red (magenta). Fluorescence images were obtained by confocal microscopy (A) and super-resolution microscopy (SR-SIM) (B). Scale bars: 10 µm (A), 1 µm (inset of A), and 1 µm (B). (C) Fluorescence images of NSUN2 (green) and BrU (magenta) were obtained by confocal microscopy. Scale bars: main, 10 µm; inset, 1 µm. (D) Fluorescence images of NSUN2 (green) and TFAM (magenta) were obtained by SR-SIM. Scale bars: main, 1 µm; inset, 0.5 µm.

### *NSUN2* KO has little impact on mitochondrial translation

NSUN2-mediated m^5^C formation stabilizes ct-tRNAs and protein synthesis (27). To determine whether mt-tRNAs are also stabilized by m^5^C modifications, we measured steady-state levels of mt-tRNAs in *Nsun2*^-/-^mouse liver by northern blotting. As reported previously (27), the level of ct-tRNA^Gly^was significantly reduced in *Nsun2*^-/-^mouse liver (Figure 5A). By contrast, we observed no significant change in the steady-state levels of mt-tRNAs upon *Nsun2*^-/-^mice (Figure 5A). Moreover, we observed no change in the steady-state level of mt-tRNAs in testis (Figure S4A) or brain (Figure S4B) of *Nsun2*^-/-^mice.

**Figure 5.**
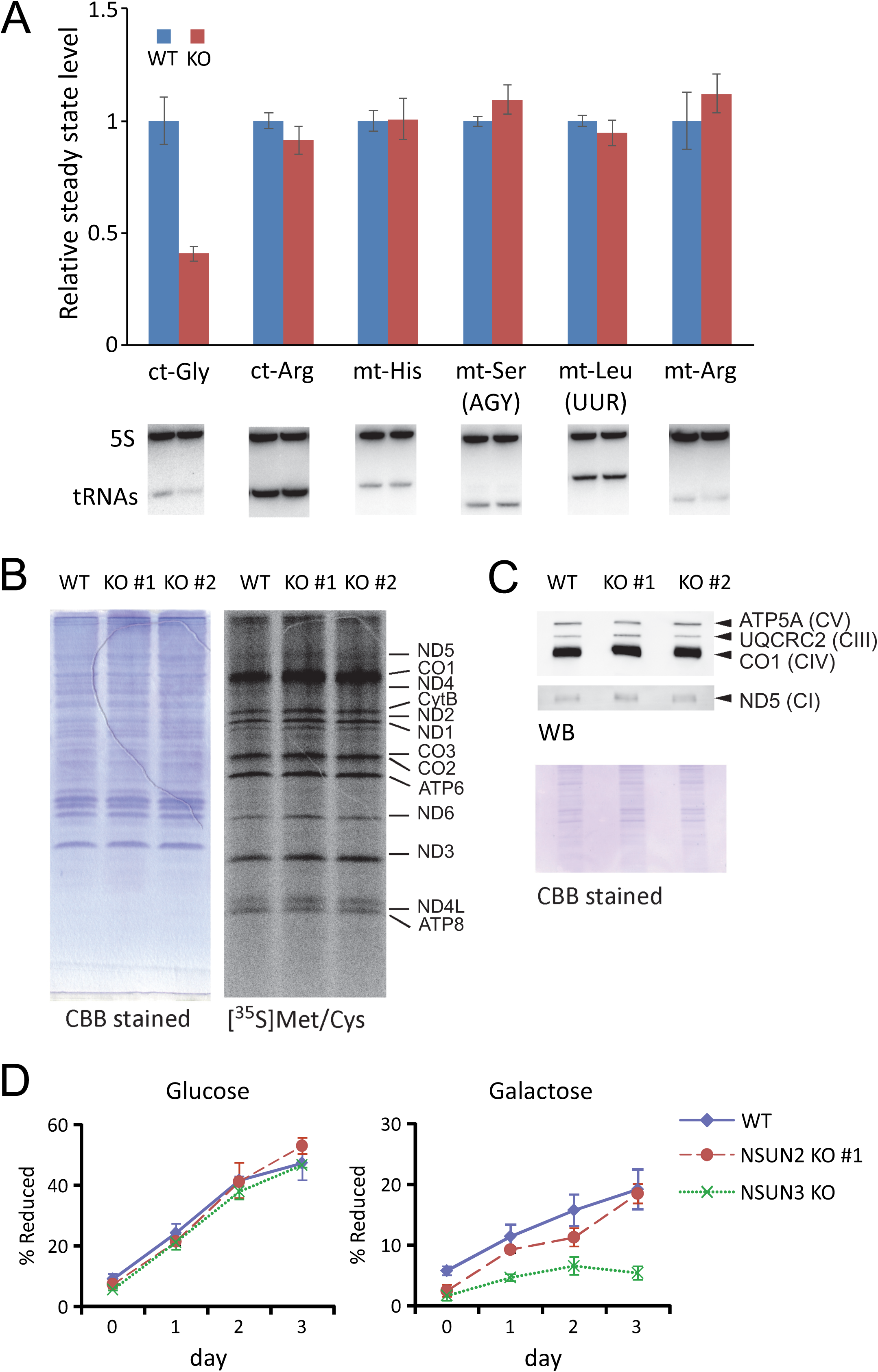
Mitochondrial translation and respiratory activity in *NSUN2* KO cells. (A) Northern blotting of ct-tRNAs and mt-tRNAs in total RNA from WT and *Nsun2*^*-/-*^ (KO) mouse liver (lower panels). Bar graph shows relative levels of each tRNA, normalized against 5S rRNA (used as a loading control). Mean ± S.D. was calculated from three biological replicates. (B) Pulse labeling of mitochondrial protein synthesis. WT and *NSUN2* KO HEK293T cells were labeled with([^35^S] Met/Cys) and chased for 1 h under emetine treatment (right). Total proteins were visualized by CBB staining (left). (C) Steady-state levels of subunit proteins in respiratory chain complexes, as determined by western blotting (top). Loaded proteins were visualized by CBB staining (bottom). (D) Growth curves of WT, *NSUN2* KO, and *NSUN3* KO HEK293T cells cultured in the medium containing glucose (left) or galactose (right) as the primary carbon source. Mean ± S.D. was calculated from three independent cultures.

Next, we performed a pulse-labeling experiment to compare mitochondrial translational activity, and observed no significant difference between WT and *NSUN2* KO HEK293T cells (Figure 5B), or WT and *Nsun2*^-/-^MEFs (Figure S5). We also analyzed steady-state levels of several subunits of the mitochondrial respiratory chain complexes by western blotting, and found no significant change upon *NSUN2* KO (Figure 5C). To explore the mitochondria-related phenotypes of *NSUN2* KO cells, we compared cell growth in galactose vs. glucose (48). As reported previously (13), growth was dramatically slowed in *NSUN3* KO cells cultured in galactose medium. *NSUN2* KO cells grew well in glucose medium, and slightly slower than WT cells in galactose medium (Figure 5D). But, this is not a significant result. Taken together, these observations indicate that *NSUN2* KO has a limited impact on mitochondrial translation and respiratory activity, notwithstanding the absence of m^5^C modification in mt-tRNAs.

## DISCUSSION

In this study, we demonstrated that NSUN2 is partially localized to mitochondria and introduces m^5^C into the extra loop of mt-tRNAs. RNA-MS detected m^5^C at eight positions in eight mouse and six human mt-tRNAs. We previously reported m^5^C at five positions in five bovine mt-tRNAs (5). Among these mammals, m^5^C is present in three species of mt-tRNAs [for Leu(UUR), Ser(AGY), and Glu], implying that m^5^C is functionally important in these tRNA species. In terms of the specific positioning of the modification, mt-tRNAs^Leu(UUR)^ and mt-tRNAs^Glu^have m^5^C48 and m^5^C49, respectively. In mt-tRNAs^Ser(AGY)^, three m^5^Cs are present at positions 48–50 in human, whereas in mouse and cow only one m^5^C is present at position 49. Even if a C is present at positions 48–50, m^5^C is not always introduced by NSUN2, e.g., C48 in mouse mt-tRNA^Leu(CUN)^ remains unmodified. Although NSUN2 is a promiscuous m^5^C methyltransferase with broad specificity for diverse RNA substrates (16), it does have some substrate and sequence preference.

Using SR-SIM, we clearly detected NSUN2 in mitochondria. In general, mitochondria-localized proteins have a mitochondria-targeting sequence (MTS) in their N-terminal region (49). However, human NSUN2 does not appear to contain an apparent MTS, and it is not described as having a mitochondrial localization in the MitoMiner database (50). Instead, human NSUN2 is predicted by WoLF PSORT to be localized to the nucleus (51). Consistent with this, we observed a strong NSUN2 signal in the nucleus of human cells (Figure 4A), as reported previously (32). However, the observation of clear foci in the mitochondria indicates that NSUN2 does in fact have a weak MTS. In further support of mitochondrial localization, NSUN2 foci co-localized with the signals of BrU-labeled RNA in mitochondria (Figure 4C), and were adjacent to the signal for TFAM (Figure 4D), a transcription factor that covers mt-DNA to form a nucleoid structure. BrU-labeled RNA in mitochondria constitutes mitochondrial RNA granules (MRGs) (52), which are sites for mitochondrial RNA processing and ribosome biogenesis, in close proximity to TFAM-stained nucleoids. Multiple factors for RNA modification and processing are localized to MRGs (53). We speculate that NSUN2 is a novel component of the MRG, where it introduces m^5^C into mt-tRNAs co-transcriptionally.

m^5^C stabilizes the RNA structure via base-pairing, base-stacking, and metal binding (54-56). Indeed, the m^5^C49 on the edge of the T-stem promotes Mg^2+^-dependent folding of tRNA (57). In canonical tRNAs, the D-loop/T-loop and D-arm/extra loop interactions are critical for the folding of tRNA into an L-shape structure (58). The D-arm/extra loop interaction is composed of a C13–G22–m^7^G46 base triple and a G15–C48 pair. G15 pairs with C48 in a reverse Watson–Crick geometry called a Levitt pair (58,59). A structural hallmark of animal mt-tRNAs is the absence of the canonical D-loop/T-loop interaction (3,60,61); instead, the D-arm and extra loop interactions, including the G15– C48 Levitt pair, are conserved in most mt-tRNAs, and play critical roles in tRNA folding and stability (3,62). In the crystal structure of tRNAs (63), m^5^C48 may strengthen the stacking interaction with a nucleobase at position 21 in the D-arm, stabilizing the mt-tRNA core structure, although the functional importance of this methylation remains to be determined.

Despite our efforts to identify a functional role of m^5^C in mt-tRNAs, we could not demonstrate any effects of this modification on mt-tRNA stability or mitochondrial protein synthesis. Further investigations will be necessary to elucidate the biological roles of m^5^C in mt-tRNAs under environmental stress conditions or in specific physiological contexts in various tissues and cells.

## ACKNOWLEDGMENTS

We are grateful to the members of the Suzuki lab, especially Yuriko Sakaguchi, Kenjkyo Miyauchi, Huan Lin and Takayuki Ohira for technical support and fruitful discussion. We also thank Drs. N. Mizushima and I. Koyama-Honda (Univ. of Tokyo) for their kind assistance with microscopic analysis. Radioisotope experiments were carried out with the support of the Isotope Science Center, The University of Tokyo. This work was supported by Grants-in-Aid for Scientific Research on Priority Areas from the Ministry of Education, Culture, Sports, Science, and Technology of Japan (MEXT) and the Japan Society for the Promotion of Science (JSPS) [26113003, 26220205, and 18H05272 to Ts.S., 26702035 to Ta.S.].

