## Supplemental information for "Mammalian NSUN2 introduces 5-methylcytidines into mitochondrial tRNAs"

A

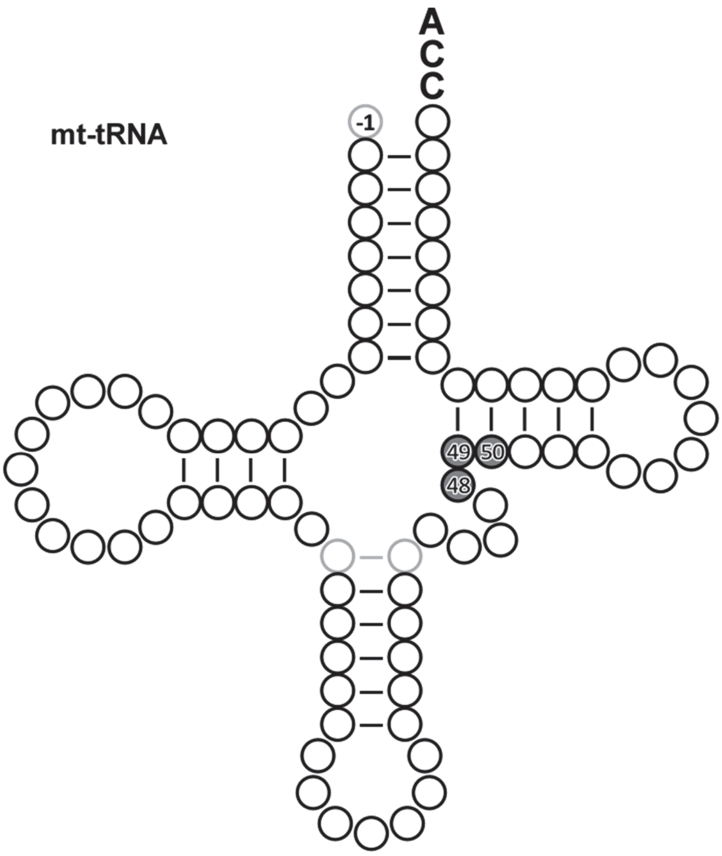


B

| | Human | Mouse | Bovine |
|---|---|---|---|
| $m^5C48$ | Phe, His, Leu(UUR), Ser(AGY), Tyr | His, Met, Leu(UUR), Asn, Tyr | Leu(UUR), Asn, Trp |
| $m^5C49$ | Glu, Ser(AGY) | Glu, Leu(CUN), Ser(AGY) | Glu, Ser(AGY) |
| $m^5C50$ | Ser(AGY) | - | - |

**Figure S1. $m^5C$ distribution in mt-tRNAs of human, mouse, and cow.**

(A) Secondary structure of mt-tRNAs. Positions of $m^5C$ modifications are indicated by shaded circles.

(B) tRNA species bearing $m^5C$ at the corresponding position in three mammals.

Bovine mt-tRNA data are from a previous study by our group (5).

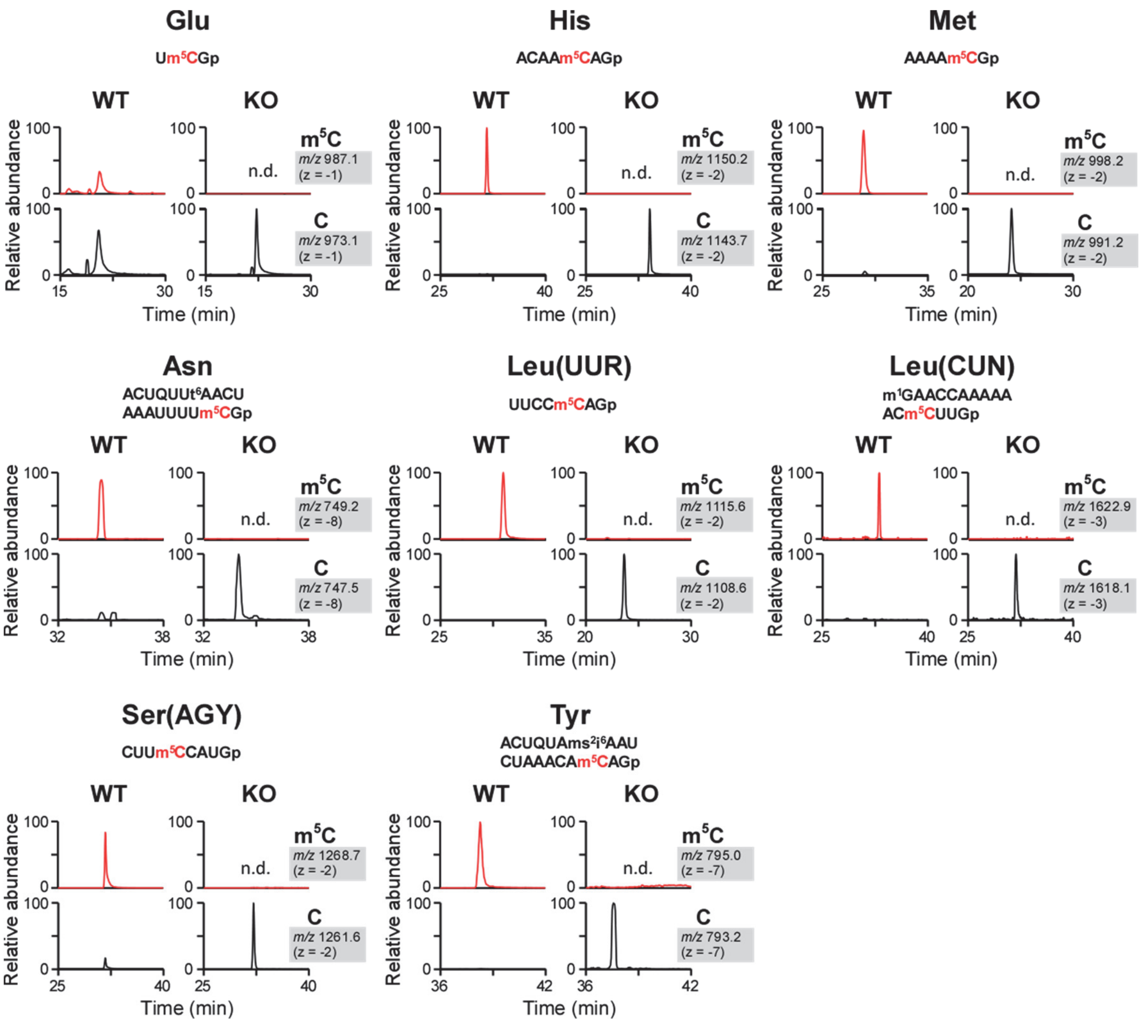


**Figure S2. m5C status in eight mt-tRNAs isolated from WT and *Nsun2* KO mouse livers**

XICs of RNase T1-digested fragments containing m5C (top) and C (bottom) in eight mt-tRNAs isolated from WT and *Nsun2*$^{-/-}$ (KO) mouse liver.

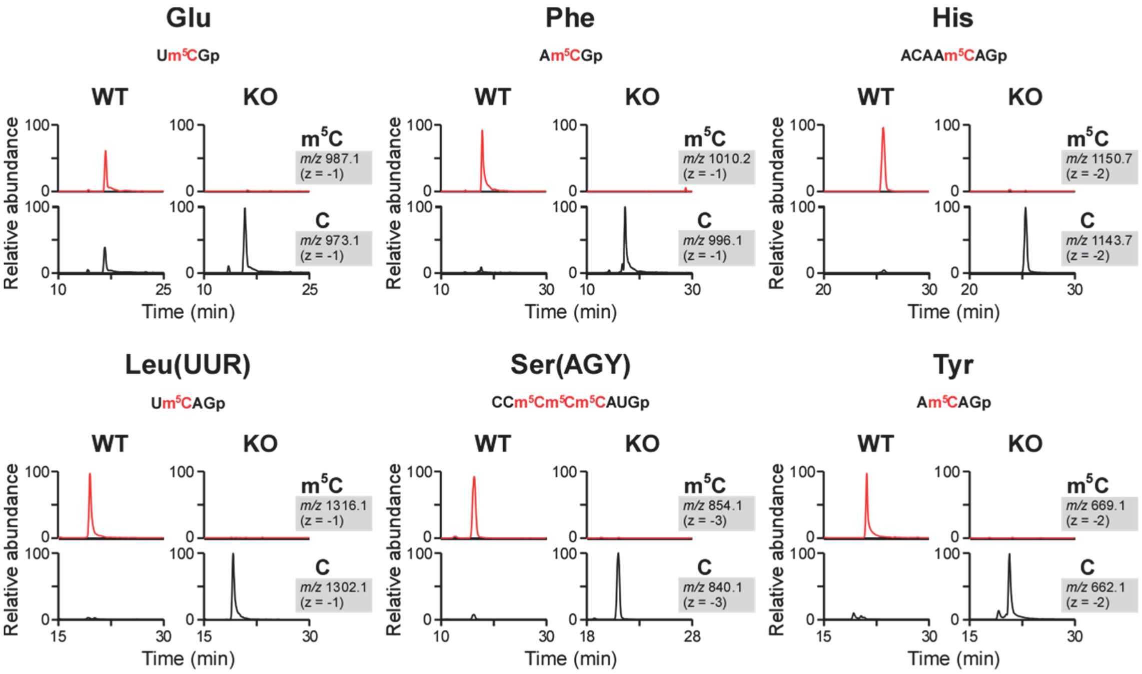


**Figure S3. m$^5$C status in six mt-tRNAs isolated from WT and *NSUN2* KO human culture cells.**

XICs of RNase T$_1$-digested fragments containing m$^5$C (top) and C (bottom) in six mt-tRNAs isolated from WT and *NSUN2* KO HEK293T cells. *m/z* value with the charge state for each fragment is shown to the right of each panel.

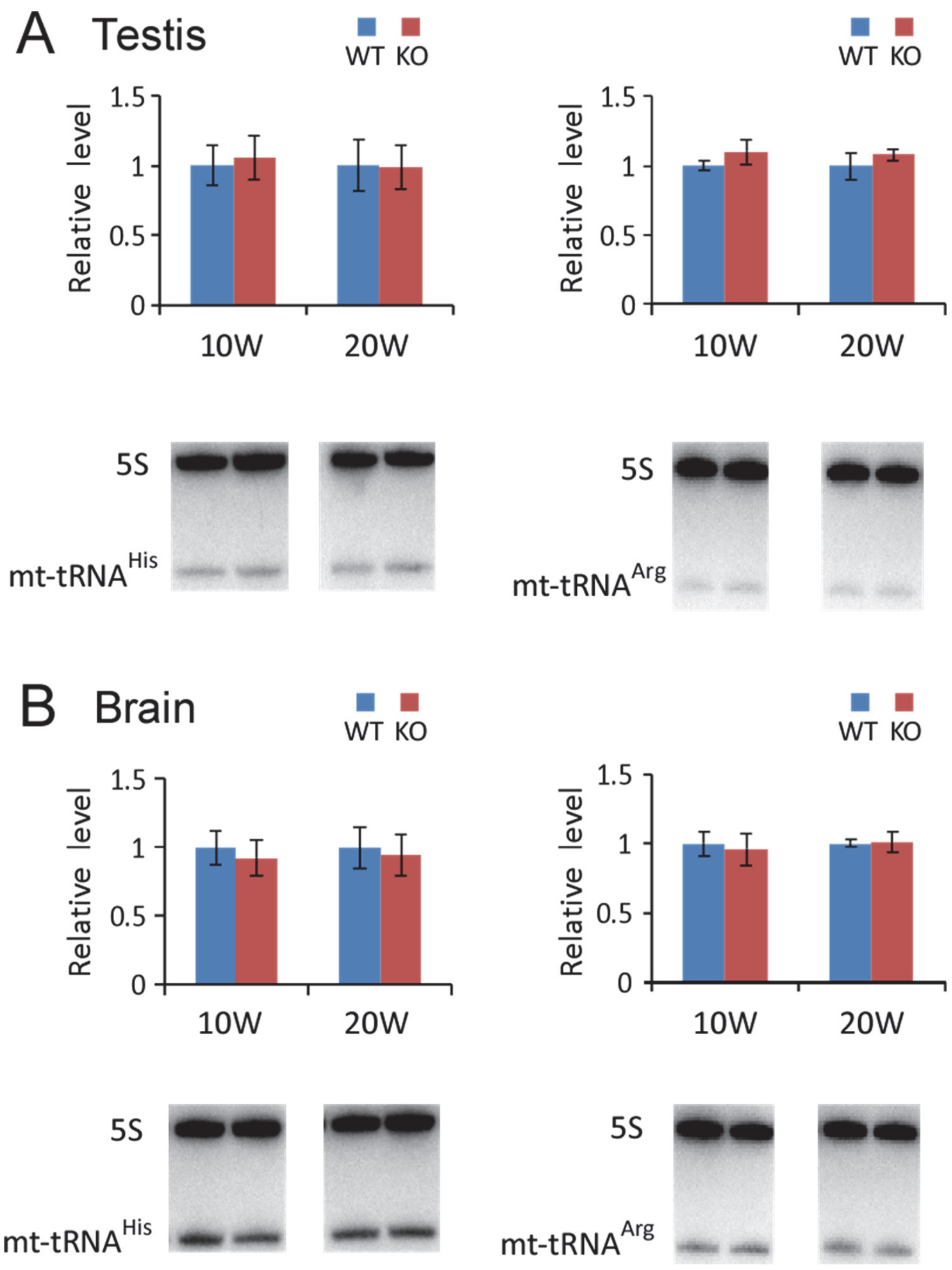


**Figure S4. Northern blotting of mouse mt-tRNAs**

Northern blots of mt-tRNAs for His (left panels) and Arg (right panels) in total RNA isolated from testis (A) and brain (B) of 10- and 20-week-old male mice (WT, blue; *Nsun2*$^{-/-}$ (KO), red). Bar graphs represent relative steady-state levels of each tRNA, normalized against 5S rRNA (used as a loading control). Means ± S.D. were calculated from three biological replicates.

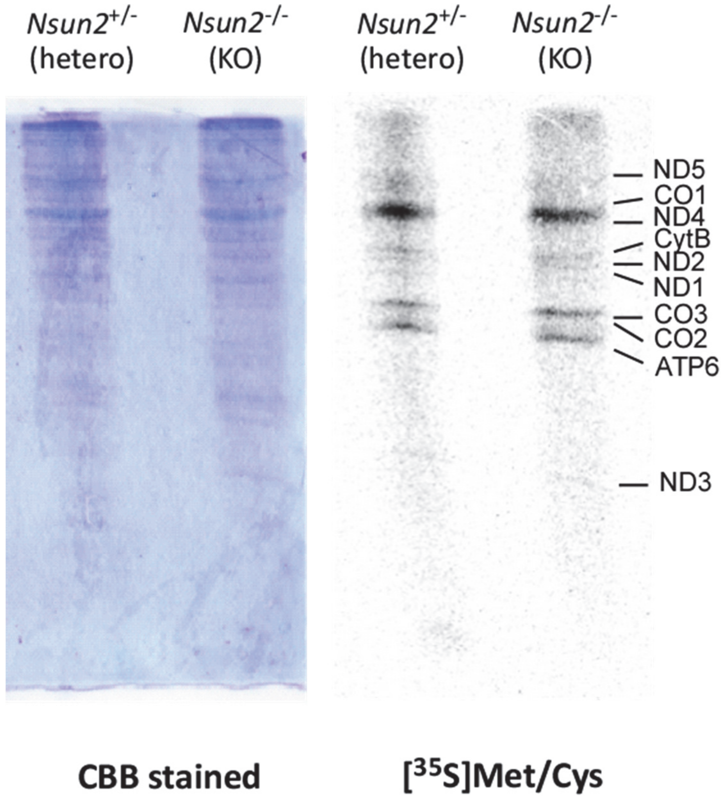


**Figure S5. Pulse labeling of mitochondrial protein synthesis in MEF cells.**
*Nsun2*$^{+/-}$ (hetero) and *Nsun2*$^{-/-}$ (KO) MEFs were labeled with ([$^{35}$S] Met/Cys) and chased for 1 h under emetine treatment (right). Total proteins were visualized by CBB staining (left).

**Table S1 List of primers and probes used in this study**

| Purpose | Name | 5' to 3' sequence |
|---|---|---|
| **Inducible mutagenesis with CRISPR/Cas9** | | |
| sgRNA for human NSUN2 gene | sgRNA_NSUN2_fw | CTTAAAGTGGCCGGGAGCG |
| | sgRNA_NSUN2_rv | ATGGCGCCGAGGGTGGTGGAAA |
| Surveyor assay | genome_NSUN2_fw2 | ACGGTTCTCTGGCACTGTAACAC |
| | genome_NSUN2_rv2 | GTGTAGGGCTAGAGTTCTGGC |
| **Isolation of mitochondrial tRNA** | | |
| For human | RCC_h-mtSer(AGY) | TGGTGAGAAAGCCATGTTGTTAGACATGGG |
| | RCC_h-mtLeu(UUR) | TATGCGATTACCGGGCTCTGCCATCTTAAC |
| | RCC_h-mtTyr | CAGTCCAATGCTTCACTCAGCCATTTTACC |
| | RCC_h-mGlu | TGGTATTCTCGCACGGACTACAACCACGAC |
| | RCC_h-mtHis | GGTAAATAAGGGGTCGTAAGCCTCTGTTGT |
| | RCC_h-mtPhe | TGGTGTTTATGGGGTGATGTGAGCCCGTCT |
| For mouse | RCC_m-mtLeu(UUR) | GAGTCTGGCGCCTTAGACCACTCGGCCATCCTGAC |
| | RCC_m-mtAsn | TTTAGTTAACAGCTAAATACCCTATTACTGGCTTCAATCT |
| | RCC_m-mGlu | ACAGCATTCAACTGCGACCAATGACATGAAAAATCATCGT |
| | RCC_m-mtHis | AAGGAGGTTTATTTCCTGTTGTCAGATTCACAGTCTAATG |
| | RCC_m-mtLeu(CUN) | TTGCACCAAGGTTTTTGGTTCCTAAGACCAATGGATTACT |
| | RCC_m-mtMet | TTCGGGGTATGGGCCCGATAGCTTAATTAGCTGAC |
| | RCC_m-mSer(AGY) | GCCATGTTTAAACATGGAAGCATGAATTAGCAGTTCTTGC |
| | RCC_m-mtTyr | AACCTCTGTGTTTAGATTTACAGTCTAATGCTTACTCAGC |
| | RCC_m-cytoGly | GAAGGGAGCTATGCTATGCTCACCACTATACCACCAACGC |
| | RCC_m-cytoLeu(CAA) | GAGTCTGGCGCCTTAGACCACTCGGCCATCCTGAC |
| **PCR primer** | | |
| Template of T7 transcription | T7 forward primer | GCTAATACGACTCACTATA |
| | mt tRNASer(AGY) WT reverse | TGGTAAGAAAGCCATGTTTAAACATGGAAGCATGAATTAGCAGTTCTTGCAATCTTTCTTTATA[illegible] |
| RT primer for cloning human NSUN2 gene | RTcDNANSUN2Rv | AGCTTGGCCAAAGAAACAAA |
| Cloning human NSUN2 gene from cDNA by nested PCR | NSUN2_1st_Fw | CCCTTAGAGCTGTTCGCTGT |
| | NSUN2_1st_Rv | CCAGAAGAAGCCAGTTTTGC |
| | NSUN2_2nd_Fw | CACCATGGGGCGGCGGTCGC |
| | NSUN2_2nd_Rv | CCACCGGGGTGGATGGACC |
| **Northern blotting** | | |
| | NT_MmmtHis_3' | TGGGGTGAATAAGGAGGTTTATTTC |
| | NT_MmmtArg | TGGTTGGTAATTATGAACAGCATCATAATC |
| | NT_MmmtLeu(UUR) | GATAGCTTAATTAGCTGACCTTACT |
| | NT_MmmtSerAGY_5' | CATGAATTAGCAGTTCTTGCAATCTTTCTT |
| | NT_Mmcyto_Gly_GCC | TCTACCACTGAACCACCCAT |
| | NT_Mmcyto_Arg_ACG | TGGCGAGCCAGCCAGGAGTCGAACCTGGAA |
| | NT_Mm_5S_rRNA | GGGTGGTATGGCCGTAGAC |
